# Coordinated overexpression of miR398 and miR408 confers broad-spectrum abiotic stress tolerance in melon

**DOI:** 10.64898/2025.12.11.693700

**Authors:** Andrea Gabriela Hernandez-Azurdia, Marta Nuñez Salvador, Javier Lozano-Ordaz, Carmelo Lopez, Ana Montserrat Martin-Hernandez, Gustavo Gómez

## Abstract

MicroRNAs (miRNAs) in general and miR398 and miR408 in particular, have emerged as key regulators of plant adaptation to individual stress, yet their roles regulating the crop response to diverse unfavorable environments remain poorly explored. Here, we present the first melon (*Cucumis melo*) plants overexpressing miR398 and miR408 precursors, two conserved miRNAs involved in the regulation of cooper (Cu) homeostasis and associated with stress-response. Engineered plants exhibited enhanced vegetative development, including stem elongation, internode formation, root architecture, and leaf production. The significant increased accumulation of well-processed miR398 and miR408, along with the downregulation of their respective targets (Copper/Zinc Superoxide Dismutase and Basic Blue Protein) demonstrating the functional activity of the transgenes. Sequencing analysis revealed positive correlation in accumulation of miR398 and miR408, suggesting a previously undescribed form of coordinated miRNA regulation, potentially independent of conventional transcription factor activity. Transgenic plants showed improved tolerance to drought, salinity, heat, and cold stress, validating the role of miR398 and miR408 as master regulators of abiotic stress resilience in melon. This work highlights the potential of biotechnological strategies based on engineering Cu-related miRNAs to enhance crop performance under different adverse environments, thereby contributing to agricultural sustainability in the face of climate change.

## INTRODUCTION

Plants are continuously exposed to a wide range of adverse environmental conditions that disrupt their normal development and productivity. In crops, these stressors often impair cellular homeostasis, resulting in substantial agricultural losses (Khan et al., 2025). Climate change, primarily driven by rising greenhouse gas emissions, is intensifying the frequency and severity of extreme weather events such as droughts, heatwaves, and floods (Bolan et al., 2024). These environmental perturbations compromise plant growth and productivity, posing a serious threat to global food security (Chaudhry and Sidhu, 2022). Current projections estimate that in 21st century, crop yields may decline by as much as 50–70% due to climate-related stressors (Rosenzweig et al., 2014; Cervantes-Godoy et al., 2014). In addition, adverse environmental conditions not only damage plants directly, but they also increase crop susceptibility to pests and diseases (Velásquez et al., 2018). Finally, the projected rise of human population to 10 billion by mid-century will require an increase of about 50% in food production compared with 2010, involving substantial expansion of agricultural lands to regions under suboptimal environmental conditions (Van Dijk et al., 2021). In light of these challenges, the development of innovative and sustainable strategies to enhance crop resilience under adverse environmental conditions is essential to safeguard agricultural yield in a changing climate scenario (Tuncel et al., 2025; Javaid et al., 2024).

Due to their sessile nature, plants have evolved sophisticated gene regulatory mechanisms to cope with environmental fluctuations and mitigate the impact of stress on physiological and cellular homeostasis. Among these mechanisms, microRNAs (miRNAs) have emerged as key post-transcriptional regulators of gene expression, playing a central role in modulating plant responses to both abiotic and biotic stresses (Shen et al., 2024; Chen et al., 2025; Liu et al., 2024).

MiRNAs are small non-coding RNAs (20 - 22 nucleotides) that constitute a critical layer of gene regulation in both animals and plants (Borges and Martienssen, 2015; Ledda et al., 2020). In plants, miRNAs can induce RNA interference at both transcriptional and post-transcriptional levels (Pradhan and Requena, 2022). They regulate gene expression in a sequence-dependent manner, influencing a wide array of biological processes, including development, environmental adaptation, and interactions with other organisms (Lee and Carroll, 2018; Pradhan and Requena, 2022).

The role of miRNAs in modulating plant responses to abiotic stress has been extensively studied in model species and major crops such as *Arabidopsis thaliana*, potato (*Solanum tuberosum*), (Lu et al., 2022b), tomato (*Solanum lycopersicum*) (Rao et al., 2022), wheat (*Triticum aestivum*) (Zhang et al., 2025), maize (*Zea mais*) (Tang et al., 2022) and rice (*Oryza sativa*) (Sun et al., 2022). In addition, it has been described that the biogenesis and turnover of certain miRNAs are events susceptible to alterations in environmental conditions (Bustamante et al., 2018; Manavella et al., 2019). However, most studies have focused on plant-response to individual stress factors (Khan et al., 2025), limiting our understanding of how miRNAs can coordinate broader regulatory networks involved in adaptation to complexes and changing environments.

To address this gap, a series of studies published between 2019 and 2021 analyzed the effects of temporal exposure to four different abiotic stressor (individually and in combination) on the miRNA population in melon (*Cucumis melo*) (Sanz-Carbonell et al., 2020, 2019; Villalba-Bermell et al., 2021). These studies identified two miRNAs, miR398 and miR408, that showed consistent and coordinated overexpression under analyzed stress conditions, suggesting a potential role in adaptive responses to a broad range of abiotic stressors. Although this hypothesis has not been experimentally validated in melon, evidence obtained unique abiotic stressors in other species such as *Medicago truncatula* (Trindade et al., 2010), maize (Qin et al., 2023), pea (*Pisum sativum*) (Jovanović et al., 2014), *A. thaliana* (Guan et al., 2013; Yang et al., 2024), rice (Yang et al., 2024), and tomato (Candar-Cakir et al., 2016; Yan et al., 2023) supports the involvement of miR398 and miR408 in regulating genes related to copper homeostasis and antioxidant defense, reinforcing their potential as global regulators of stress resilience (Li et al., 2022; Ferreira-Silva et al., 2025; Yang et al., 2024; Li et al., 2024b). Additionally, miR398 and miR408 represent two of the scare miRNA-families that are deeply conserved in land plants (Cuperus et al., 2011; Guo et al., 2020), indicating that they play a key role in the regulation of the plant adaption to terrestrial habitats (Cuperus et al., 2011).

Melon is an economically important crop, widely cultivated in semi-arid regions across the globe (Wei et al., 2023), and particularly sensitive to environmental stress, especially in Mediterranean areas where heatwaves and water scarcity directly affect yield (López-Sesé, 2023). Its economic relevance, combined with the availability of genomic resources such as a high-quality reference genome (Garcia-Mas et al., 2012; Zhang et al., 2019) and functional tools, makes melon a suitable model for studying miRNA regulatory networks in crops. Moreover, its well-characterized genetic diversity enables the design of targeted improvement strategies for specific cultivars. However, melon is considered a recalcitrant species for stable genetic transformation, which has resulted in a significantly lower number of transgenic lines compared to other major crops, such as tomato (Shirazi Parsa et al., 2023). This limitation has slowed the functional validation of candidate genes and regulatory elements, including miRNAs, in melon, underscoring the need for optimized transformation protocols and alternative biotechnological approaches to accelerate genetic improvement in this species.

Manipulating endogenous miRNAs is emerging as a promising biotechnological strategy to enhance plant resilience to environmental stress, offering the ability to reconfigure stress response networks in a targeted and efficient manner (Ferreira-Silva et al., 2025; Basso et al., 2019). In this context, the term *targeted* describes the specificity of a genome-modifying strategy in recognizing/interacting with a particular DNA/RNA sequence (Li et al., 2024a). Compared to conventional transgenesis or gene editing, miRNA-based approaches allow for finer and reversible regulation of stress-related genes, facilitating crop adaptation without compromising performance under optimal conditions (Shang et al., 2023).

In this study, we present the first transgenic melon plants (*Cucumis melo* var. cantaloupe, cv. Vedrantais) in that the biogenesis of miRNAs was modified. Two diploid lines constitutively overexpressing miR398a and miR408 precursors were generated via stable transformation using *A. tumefaciens* and characterized at phenotypic, transcriptomic, and small RNA levels under both control and abiotic stress conditions (drought, salinity, cold, and heat). The modified plants exhibited enhanced tolerance to the four adverse environmental conditions across various vegetative stages, supporting the hypothesis that regulating miR398 and/or miR408 expression is a promising strategy to improve global abiotic stress resilience in melon.

In summary, this work represents a pioneering effort in the functional validation of the role of miRNAs as regulators of the stress-response in melon, providing experimental evidence for the coordinated action of miR398 and miR408 under abiotic stress, and lays the foundation for innovative biotechnological strategies aimed to improve yield to address global food security and developing sustainable agricultural practices in the context of global climate change.

## RESULTS

### Generation of functional analyses of transgenic lines

To study the role of miR398a and miR408 in melon development and its response to abiotic stress conditions, constructs containing the precursors of these miRNAs (Fig. 1A) under the control of the 2x35S promoter and the Pop-it terminator were generated. The precursors were cloned into the binary vector pMCD32 using BsaI restriction sites, which originally contained the lethal gene ccdB and a chloramphenicol resistance cassette (CmR). Insertion of the precursors replaced this cassette allowing the identification of the positive constructions. Both constructs include the basta resistance gene (bar), regulated by the NOS promoter and terminator, as well as the CaMV polyadenylation signal (Fig. 1B).

**Figure 1.**
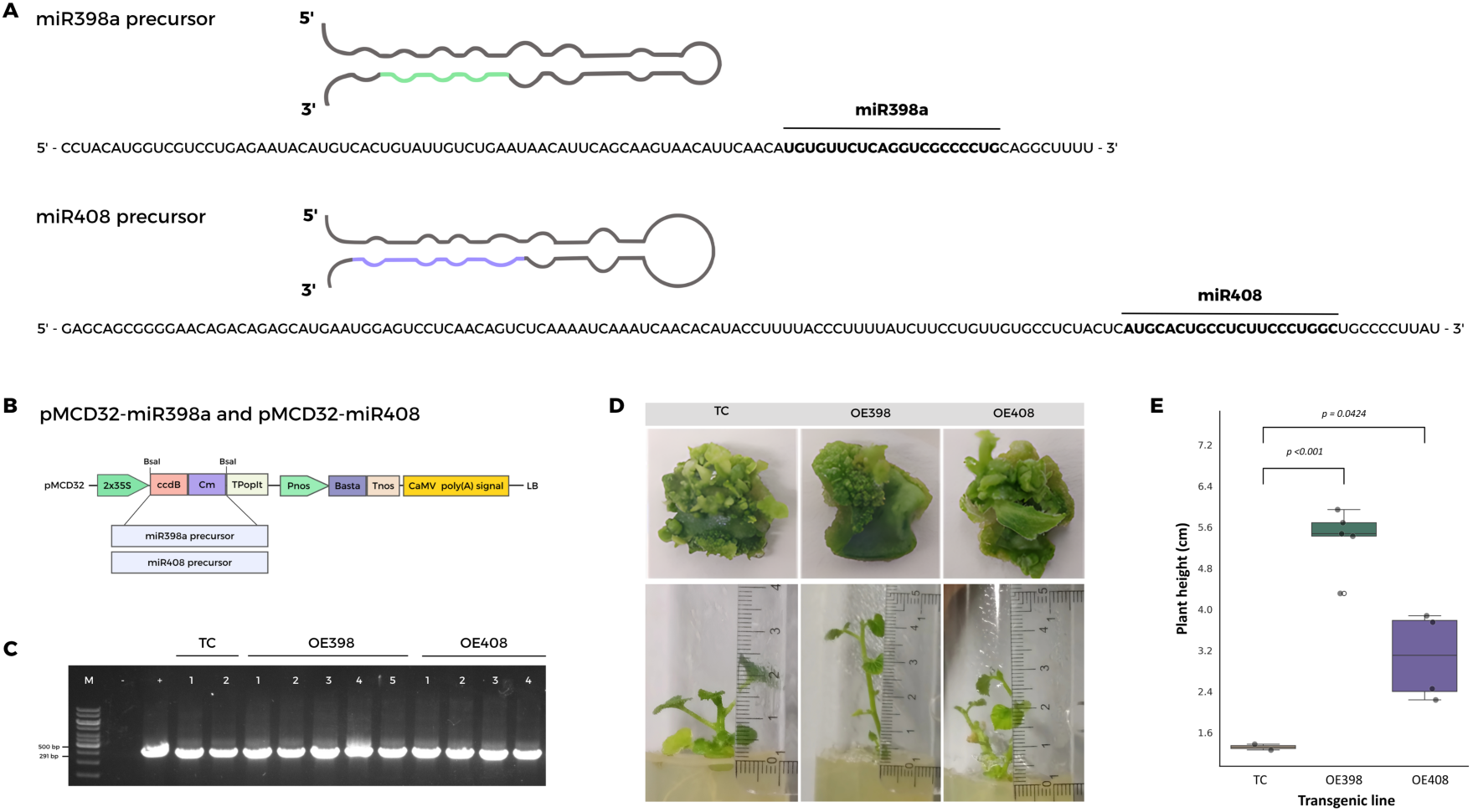
Molecular and morphological characterization of transgenic melon explants overexpressing miR398a and miR408 precursors. (A) Predicted secondary structures and nucleotide sequences of the miR398a and miR408 precursors in melon, with mature sequences highlighted. (B) Schematic representation of the binary vectors pMCD32-miR398a and pMCD32-miR408, containing the precursors under the control of the 2×35S promoter and the Pop-it terminator. Both vectors include the BASTA resistance gene (bar), regulated by the NOS promoter and terminator, and the CaMV polyadenylation signal. The precursors were inserted into the BsaI restriction sites, replacing the ccdB/CmR negative selection cassette. (C) Gel electrophoresis of representative PCR products confirming construct integration by amplification of the bar gene (291 bp) in the transgenic lines. (D) Representative images of explants cultured for 24 days in regeneration medium, showing morphological differences between the TC (transformed control), OE398, and OE408 lines. Representative individualized plants derived from the explants are also shown. (E) Boxplot showing the significant differences in length between plants derived from the TC and OE398/OE408 explants.

A total of 200 explants per construct were initially used for transformation. Transformed explants were selected based on their resistance to glufosinate, conferred by the bar gene, by culturing them on regeneration medium supplemented with the herbicide. Notable differences in the regenerative capacity of the miR398/miR408 and control explants were observed between the different lines evaluated. After 24 days of culture in regeneration medium, explants from lines transformed with miR398a (OE398) and miR408 (OE408) precursors showed more abundant development of meristematic structures and regenerative tissue compared to explants from the transformed control (TC). In particular, OE398 explants displayed a high density of compact green structures, while OE408 explants showed extensive areas of less compact but well-developed regenerative tissue. In contrast, TC explants exhibited a reduced number of regenerative structures and limited growth (Fig. 1D, upper panel). The integration of the constructs in explant-derived plants, was confirmed by PCR using primers specific for the bar gene (Bar-F and Bar-R), based in the amplification of the expected 291-bp fragment indicative of positive transformation (Fig. 1C).

Consistently, explant-derived individualized plants showed differences in vegetative growth (Fig. 1D, lower panel). After 15 days of culture on rooting medium OE398 and OE408 plants reached medium heights of 5.28 and 3.08 cm, respectively. While TC plants were significantly smaller, with average heights of 1.31 cm. A total of 15, 12 and 2 plants derived from OE398, OE408 and TC explants, respectively were analyzed to confirm the increased expression of the corresponding miRNA-precursors (Fig. Supplementary 1). Positively transformed lines were then analyzed by flow cytometry to determine their ploidy status. After this selection steps, we selected for homozygous generation and subsequent analysis a total of five diploid lines for OE398, four for OE398 and one for TC constructs, respectively (marked in bold in the Fig. Supplementary 1).

To evaluate the correct processing and functional activity of miRNA-precursor transgenes, expression analyses were conducted to quantify the accumulation of miR398a and its target gene Copper/Zinc Superoxide Dismutase (CSD), as well as miR408 and its target gene Basic Blue Protein (BBP). Transgenic melon plants derived from five independent OE398 lines (named OE398-1 to OE398-5) and four independent OE408 lines (named OE408-1 to OE408-4) were evaluated.

Quantitative stem-loop RT-qPCR analysis revealed a consistent increase in the accumulation of miR398a and miR408 across all transgenic lines compared to the transformed control (Fig. 2 and Table S1). In parallel, significant downregulation of their respective target transcripts (CSD and BBP) was observed, indicating effective regulation mediated by the overexpressed miRNAs. Among the analyzed lines, OE398-4 and OE408-1 showed the highest levels of miRNA accumulation, which correlated with the lowest expression levels of their respective target genes. Additionally we confirmed an increased accumulation of both miR398a and miR408 precursors in these transgenic lines (Fig. Supplementary 2). Based on these results, lines OE398-4 and OE408-1 were selected for further phenotypic characterization and stress tolerance assays.

**Figure 2.**
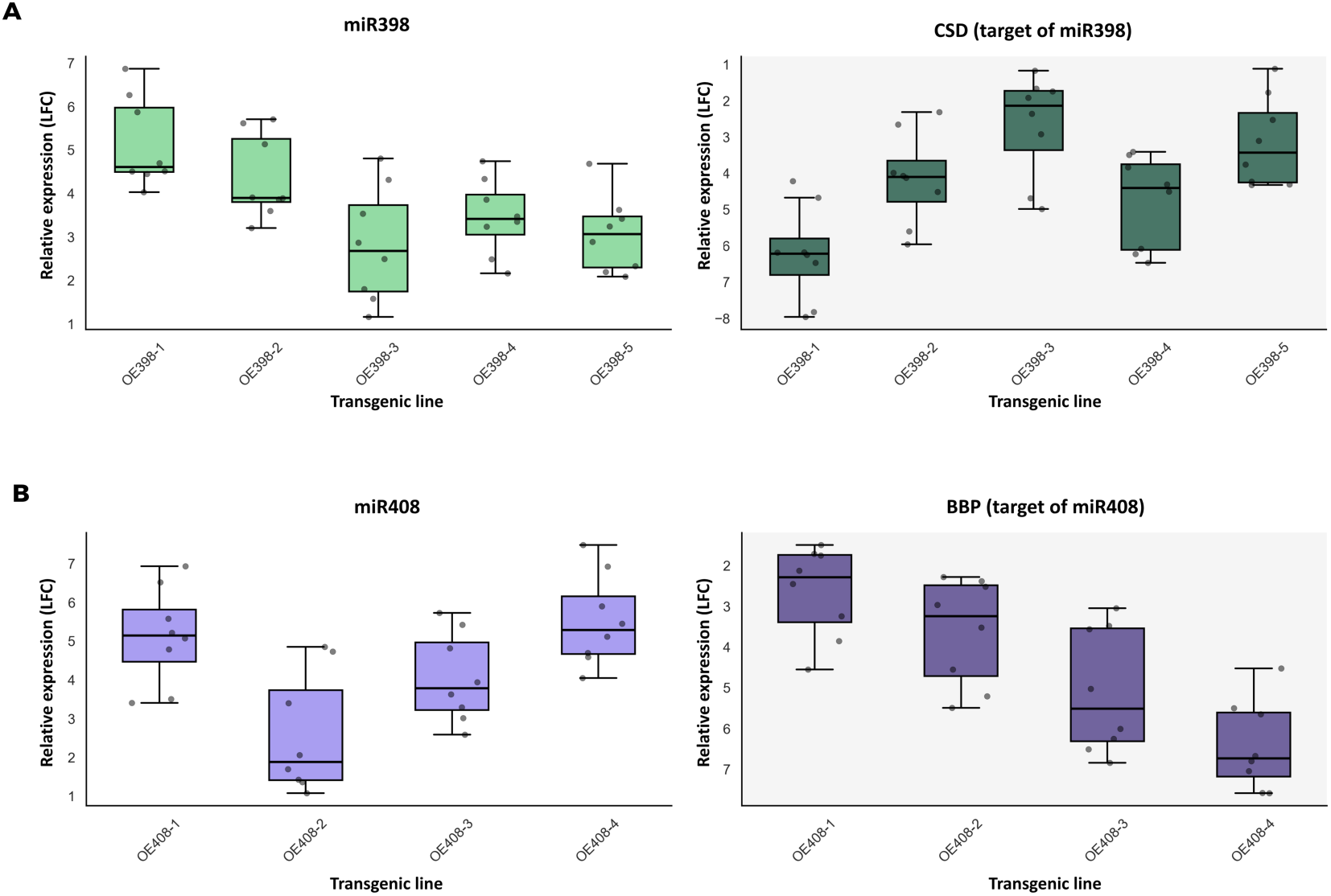
Expression analysis of miR398 and miR408 and their target genes in transgenic melon lines. (A) Relative expression levels of miR398 (left) and its target gene Copper/Zinc Superoxide Dismutase (CSD, right) in five independent OE398 lines. (B) Relative expression levels of miR408 (left) and its target gene Basic Blue Protein (BBP, right) in four independent OE408 lines. Box plots display the distribution for each line, with individual data points superimposed.

### OE398-1 and OE408-4 lines exhibit enhanced vegetative development

Prior to the detailed characterization of the transgenic lines, a comparative analysis was conducted between non-transformed and TC melon plants, focusing on morphological traits and miR398/miR408 expression levels. The results indicated that both non-transformed and TC plants showed comparable vegetative development, with no evident differences in size and overall architecture (Fig. Supplementary 3A). As expected, expression levels of miRNA precursors was comparable in both untransformed and TC plants, with no significant variation in miR398 or miR408 precursor accumulation (Fig. Supplementary 3B). Having confirmed the absence of significant developmental differences between TC and non-transformed plants and that transformation *per se* does not alter basal expression of the miRNA precursors, we proceeded to evaluate the effects of miR398a and miR408 overexpression in OE398-1 and OE408-4 transgenic lines, using plants of TC line as reference.

Although all plants were maintained under controlled conditions for 80 days after germination (DAG), phenotypic analyses were performed at 30 DAG when miRNA overexpressing plants showed evident differences with TC (Fig. 3A and 3B). At this stage, OE398-1 and OE408-4 plants exhibited marked phenotypic differences compared to TC controls, including increased size, enhanced foliage development, and greater overall vigor (Fig. 3A). Stem elongation was significantly higher in the transgenic lines, with average heights of 42.4 cm (OE398-1) and 31.8 cm (OE408-4), compared to 15.3 cm in TC plants (Fig. 3B). These differences remained consistent throughout the 80-day evaluation period.

**Figure 3.**
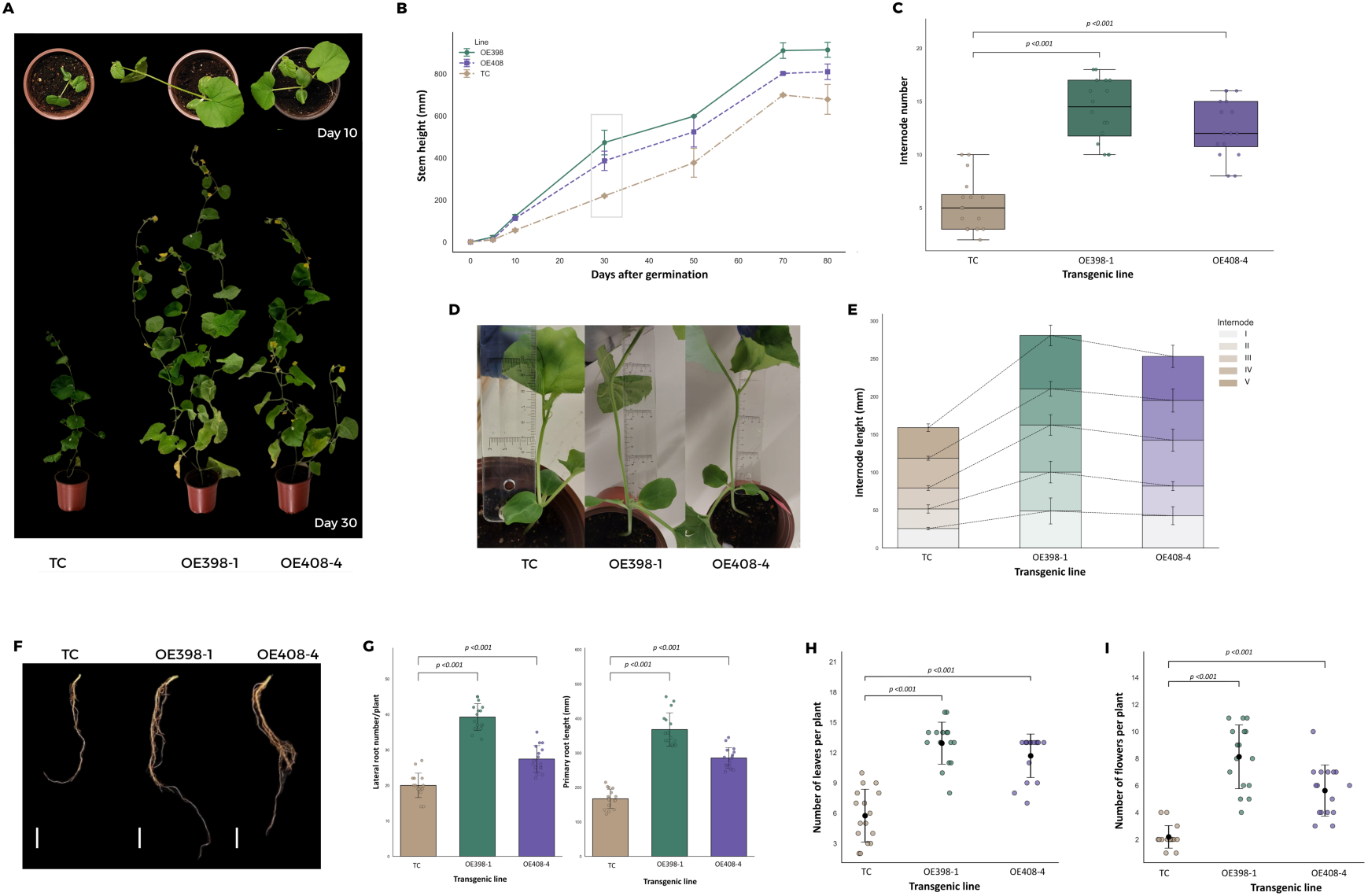
Morphological evaluation of transgenic melon lines OE398 and OE408. (A) Representative images of TC, OE398, and OE408 plants at 10 and 30 days after germination (DAG), showing visible differences in size, leaf density, and overall vigor. (B) Growth curves indicating significantly higher elongation rates in OE398 and OE408 compared to TC. At 30 DAG, OE398 and OE408 reached average heights of 42.4 cm and 31.8 cm, respectively, while TC averaged 15.3 cm (p < 0.01; n.s. between OE398 and OE408). Subsequent measurements were performed at 30 DAG. (C) Number of internodes per plant, with OE398 and OE408 averaging 14.2 and 13.0, respectively, versus 4.1 in TC. (D) Representative image illustrating the procedure for measuring the first internode length in TC, OE398, and OE408 plants. (E) Cumulative length of the first five internodes, significantly greater in OE398 (22.7 cm) and OE408 (22.3 cm) compared to TC (15.4 cm). (F) Representative images of root systems, showing increased lateral root density and length in OE lines. (G) Root length, with OE398 and OE408 averaging 39.3 cm and 30.9 cm, respectively, versus 18.1 cm in TC (p < 0.01). (H) Number of leaves per plant, significantly higher in OE398 (13.2) and OE408 (12.8) compared to TC (5.9) (p < 0.01). (I) Flower production per plant, with OE398 and OE408 averaging 8.3 and 5.0 flowers, respectively, compared to 2.1 in TC (p < 0.05).

Internode analysis revealed a significant increase in the number of internodes in OE398-1 and OE408-4 plants, with averages of 14.2 and 13.0 internodes per plant, respectively, versus 4.1 in TC plants (Fig. 3C). Additionally, the cumulative length of the first five internodes was significantly greater in the transgenic lines, reaching 22.7 cm in OE398-1 and 22.3 cm in OE408-4, compared to 15.4 cm in TC plants (Fig. 3D and 3E).

Root system development was also enhanced in the transgenic lines (Fig. 3F). Root length measurements showed averages of 39.3 cm (OE398-1), 30.9 cm (OE408-4), and 18.1 cm (TC) (Fig. 3G). Similarly, the number of lateral roots was significantly higher in OE398-1 and OE408-4 plants, with averages of 37.2 and 32.8, respectively, compared to 20.3 in TC plants.

Leaf production was also increased in both miRNA-overexpressing lines. As shown in Figure 3H, OE398-1 and OE408-4 plants produced an average of 13.2 and 12.8 leaves per plant, respectively, compared to 5.9 in TC controls. Flower production also improved, with OE398-1 and OE408-4 plants generating an average of 8.3 and 5.0 flowers per plant, respectively, versus 2.1 in TC plants (Fig. 3I).

Taken together, these results demonstrate that overexpression of miR398 and miR408 in melon leads to a consistent and significant enhancement of vegetative development, including stem elongation, internode formation, root architecture, leaf production, biomass accumulation, and flowering capacity.

### miR398 and miR408 Overexpression Induces Modifications in Both sRNA and Coding Transcript Profiles

To determine the precise processing of both miRNA-precursors and assess the global impact of miR398 and miR408 overexpression in OE398-1 and OE408-4 lines, we performed a comprehensive transcriptional analysis at both the small RNA (sRNA) and coding transcript levels. A total of 12 libraries (six for total RNA-seq and six for small RNA-seq), including two biological replicates per transgenic line, were generated. Detailed information regarding raw and trimmed read count and sequences quality for each omic approach is provided in Table S2.

Global transcriptional profiles (including TC, OE398-1, and OE408-4 plants and their biological replicates) were evaluated using Principal Component Analysis (PCA). The first two principal components explained 82.0% of the total variance for sRNAs and 89.0% for coding transcripts. In both analyses, according to the results of the Mann-Whitney-Wilcoxon test, the distance between groups was significantly greater (p < 0.05) than the intra-group distance, indicating that most of the transcriptional variation can be associated to the transgenic condition and supporting the robustness of the experimental data (Supplementary Fig. 4).

The accumulation profile of endogenous sRNAs (ranging from 20- to 25-nt) across the three analyzed lines was consistent with previously reported data for melon cultivars (Sanz-Carbonell et al., 2019), showing a predominance of 24-nt sRNAs (Fig. 4A and table S3). As determined by a multivariate permutational analysis (PERMANOVA, *p* = 0.133), no significant differences in sRNA size distribution were observed among the three analyzed lines. Based on the parameters described in the Materials and Methods section, we determined the number of endogenous sRNAs that showed significantly altered accumulation in the OE398-1 and OE408-4 lines (1174 and 636 unique reads, respectively) compared to TC plants (Fig. 4B and 4C). Upregulation was the predominant response, with 935 and 574 unique reads upregulated in OE398-1 and OE408-4, respectively, while 239 and 62 unique sequences were downregulated. A total of 458 unique sRNA sequences showed common differential expression in both OE398-1 and OE408-4 lines (Fig. 4D and Table S4). Common differentially expressed sRNAs were predominantly derived from tRNAs and intronic regions (Fig 4E and Table S4). Spearman correlation analysis using Log₂ Fold Change values of the common reactive sRNAs revealed a significant positive correlation (ρ = 0.852; p-value = 2.37 × 10^-30^) between the profiles of the two lines (Fig. 4F and Table S5), indicating a strong correspondence in the reprogramming of shared differential sRNAs.

**Figure 4:**
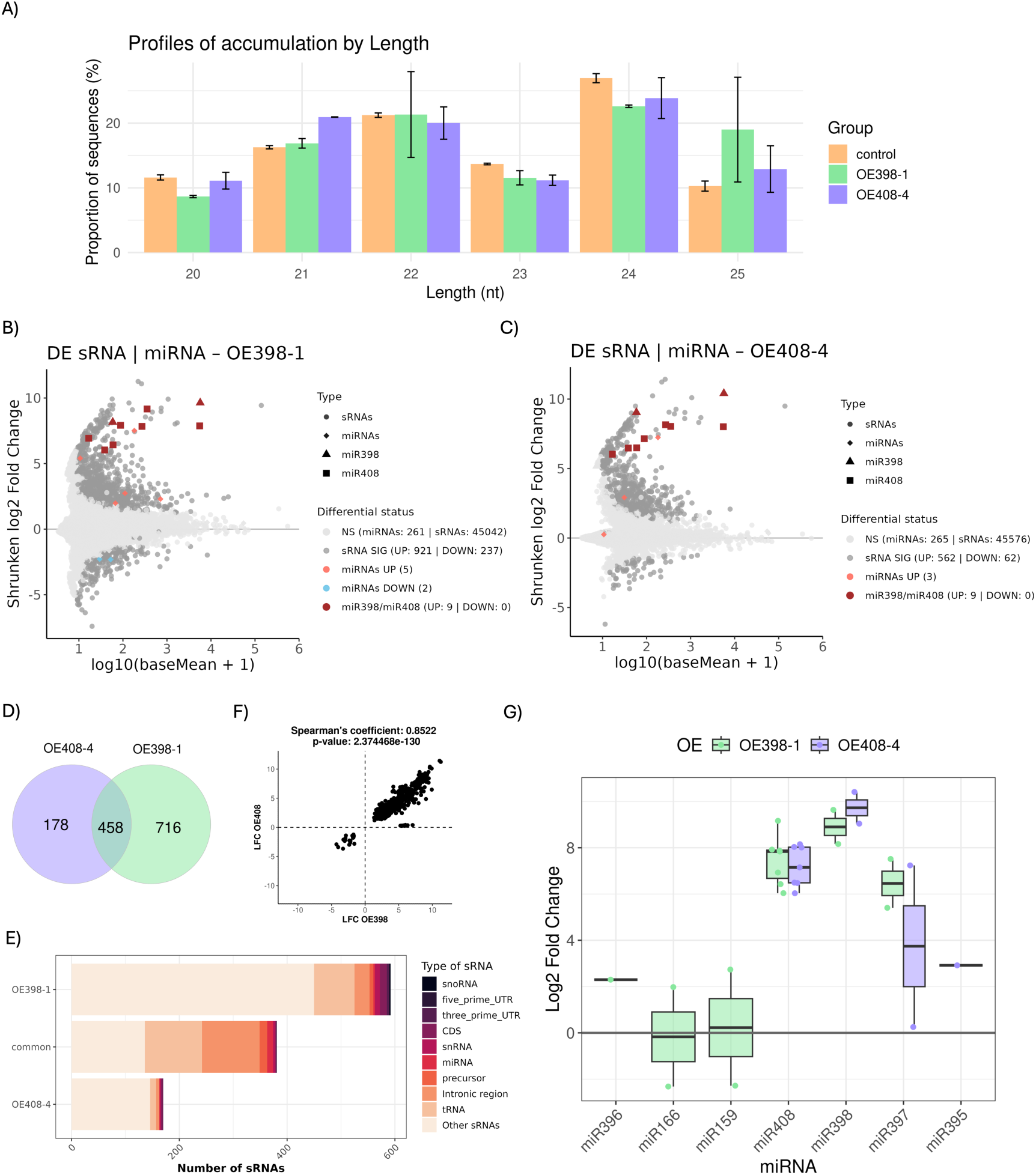
Global analysis of the sRNA population in OE398-1 and OE408-4 plants. **A**) Diagram showing the means of the relative accumulation and distribution of the total sRNAs reads ranging 20 to 25 nt (Error bars indicate the SE). The three analyzed lines are represented by colors (orange: TC, green: OE398-1, and purple: OE408-4). MA plot representing melon sRNAs with significant differential expression in OE398-1 (B) and OE408-4 (C) plants. Dark red triangles and squares represent miR398-related and miR408-related sequences, respectively. The number of differential sRNAs in each transgenic line is detailed. **D**) Venn diagram representing the number of differential sRNAs. E) Graphic representation of the type of differential sRNAs recovered from OE398-1 and OE408-4 plants. F) Graphic showing the correlation in the expression of sRNAs identified as differential in both OE398-1 and OE408-4 plants. **G**) Box-plot analysis showing the general expression value observed for each miRNA-family member (dots) in OE398-1 (green) and OE408-4 (purple) plants. The differential expression of each miRNA family is represented by the median (internal box line) of the LFC values.

Regarding the specific accumulation of miRNAs derived from the overexpressed precursors (pre-miR398a and pre-miR408), we observed (consistent with stem-loop RT-qPCR results) that OE398-1 and OE408-4 lines exhibited increased levels of sequences corresponding to miR398a (LFC= 8.90) and miR408 (LFC= 7.15), respectively (Fig. 4G). These highly accumulated sequences correspond predominantly to the canonical forms of mature miR398 and miR408, indicating that the overexpressed precursors were accurately processed in the transgenic OE398-1 and OE408-4 plants (Fig. Supplementary 5). A broader analysis of melon miRNA families revealed that overexpression of miR398 and miR408 precursors led to slight alterations in the global miRNA profile of the transgenic plants. Members of five miRNA families (miR408, miR397, miR159, miR396, and miR166) showed differential accumulation in OE398-1, while three miRNAs (miR398, miR397, and miR395) were affected in OE408-4 (Fig. 4F). Except for miR166d and miR159b (in OE398-1 plants) the remains miRNAs showed significantly increased accumulation in the two transgenic lines (Fig. 4G and Table S6).

Three miRNAs (miR398a, miR408, and miR397) exhibited shared differential expression in both OE398-1 and OE408-4 plants (Fig. 4G and table S6). The most notable overlap in accumulation levels was observed between miR398a and miR408. Expression analysis revealed that in OE408-4 plants, the overexpression of miR398a (LFC=9.73) was comparable to that of miR408, consistent with the observed for the increased accumulation of miR408 (LFC=7.84) in OE398-1 plants. Differential accumulation of LFC=6.46 and LFC=3.74 for miR397 was observed in OE398-1 and OE408-4 lines.

Considering the transcriptional landscape associated with OE398-1 and OE408-4 plants, we identified 836 unique differentially expressed transcripts in OE398-1 (Fig. 5A) and 414 in OE408-4 lines (Fig. 5B). Unlike the pattern observed for sRNAs, the number of up- and down-regulated transcripts was comparable in both lines. Specifically, 449 transcripts were upregulated and 387 downregulated in OE398-1, while 224 were upregulated and 190 downregulated in OE408-4 plants.

**Figure 5:**
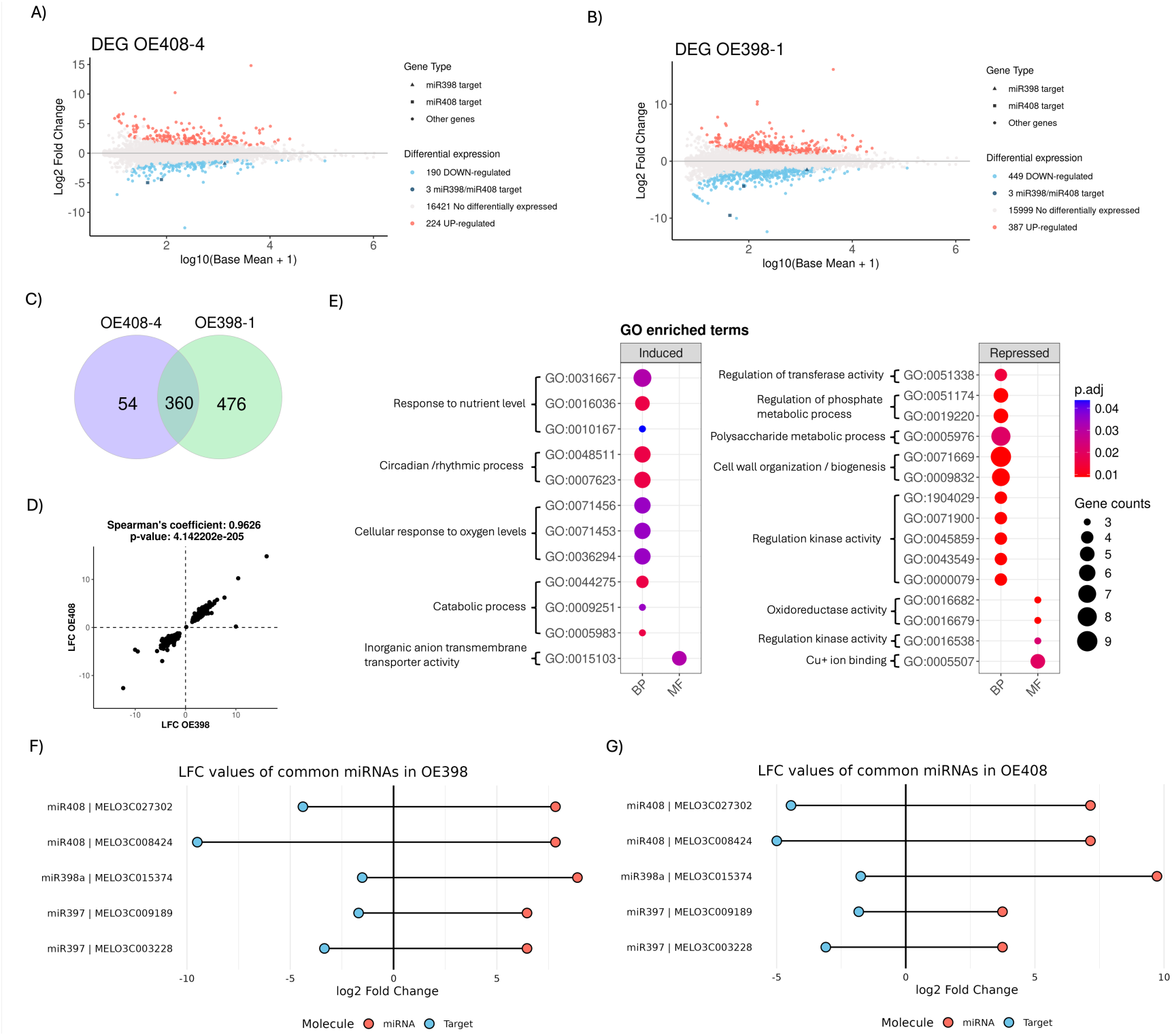
Transcriptional alterations associated in OE398-1 and OE408-4 plants. MA plot representing diferentially expressed genes (DEG) in OE398-1 (A) and OE408-4 (B) plants. Blue and red dots indicate significant up- and down-regulated genes, respectively. Dark blue triangles and squares represent miR398- and miR408-targets, respectively. The number of DEG in each transgenic line is detailed. C) Venn diagram representing the number of DEG. D) Graphic showing the correlation in the expression of transcripts identified as differential in both OE398-1 and OE408-4 plants. **E**) Gene ontology analysis for DEGs shared in both OE398-1 and OE408-4 plants. Circle size represents the number of DEG associated to the GO term (lateral scale). Color indicates p. adj values. BP: Biological process, MF: Molecular funtion. Graphic representation of relative accumualtion of common miRNAs (red dots) overexpressed in both OE398-1 (F) and OE408-4 (G) plants and their respective targets (blue dots).

A total of 360 transcripts showed common differential expression in both transgenic lines (Fig. 5C and Table S7). To quantify this shared transcriptional response, we performed a Spearman correlation analysis using Log₂ Fold Change values of the commonly regulated transcripts in OE398-1 and OE408-4 plants. The results revealed a significant positive correlation (ρ = 0.963; p-value = 4.14 × 10⁻²⁰⁵) between the transcriptional profiles of the two lines, indicating a strong correspondence in their transcriptional reprogramming (Fig. 5D). Gene Ontology (GO) term analysis of the common transcriptional response to miR398 and miR408 overexpression revealed that, in general, genes related to responses to external factors and circadian processes were predominantly repressed in OE398-1 and OE408-4 plants (Fig. 5E and Table S8). Meanwhile, genes involved in the regulation of kinase and transferase activity, metabolic processes, and cell wall biogenesis were repressed in both transgenic lines. Considering enriched molecular functions, a marked decrease in Cu⁺ binding activity was evident, along with an increase in transmembrane transport.

Regarding melon transcripts targets of the commonly overexpressed miRNAs in OE398-1 (Fig. 5F and table S9) and OE408-4 (Fig. 5G and Table S9) plants, transcriptional data revealed that CSD (target of miR398), BBP (target of miR408), and Laccase-related genes (targets of miR397) were significantly downregulated in both transgenic lines, supporting the functional activity of these overaccumulated miRNAs.

### OE398-1 and OE408-4 plants exhibit enhanced tolerance to diverse abiotic stress conditions

To determine whether overexpression of miR398a and miR408 affects plant responses to adverse environmental conditions, we monitored growth patterns in OE398-1 and OE408-4 transgenic melon lines exposed for 8 days to four types of abiotic stress: heat, drought, cold, and salinity. The 8-day stress exposure period was established based on the survival capacity of TC plants under heat, cold, and salinity conditions.

Overall, OE398-1 and OE408-4 plants exhibited more vigorous growth than the control line (TC), showing clear greater height and fresh weight, along with reduced wilting particularly under heat and drought stress (Fig. 6A). Under cold and salinity conditions, OE398-1 and OE408-4 plants maintained a more compact and healthy architecture, whereas TC plants displayed clear signs of stress, including chlorosis, reduced biomass, and impaired leaf development (Fig. 6A).

**Figure 6.**
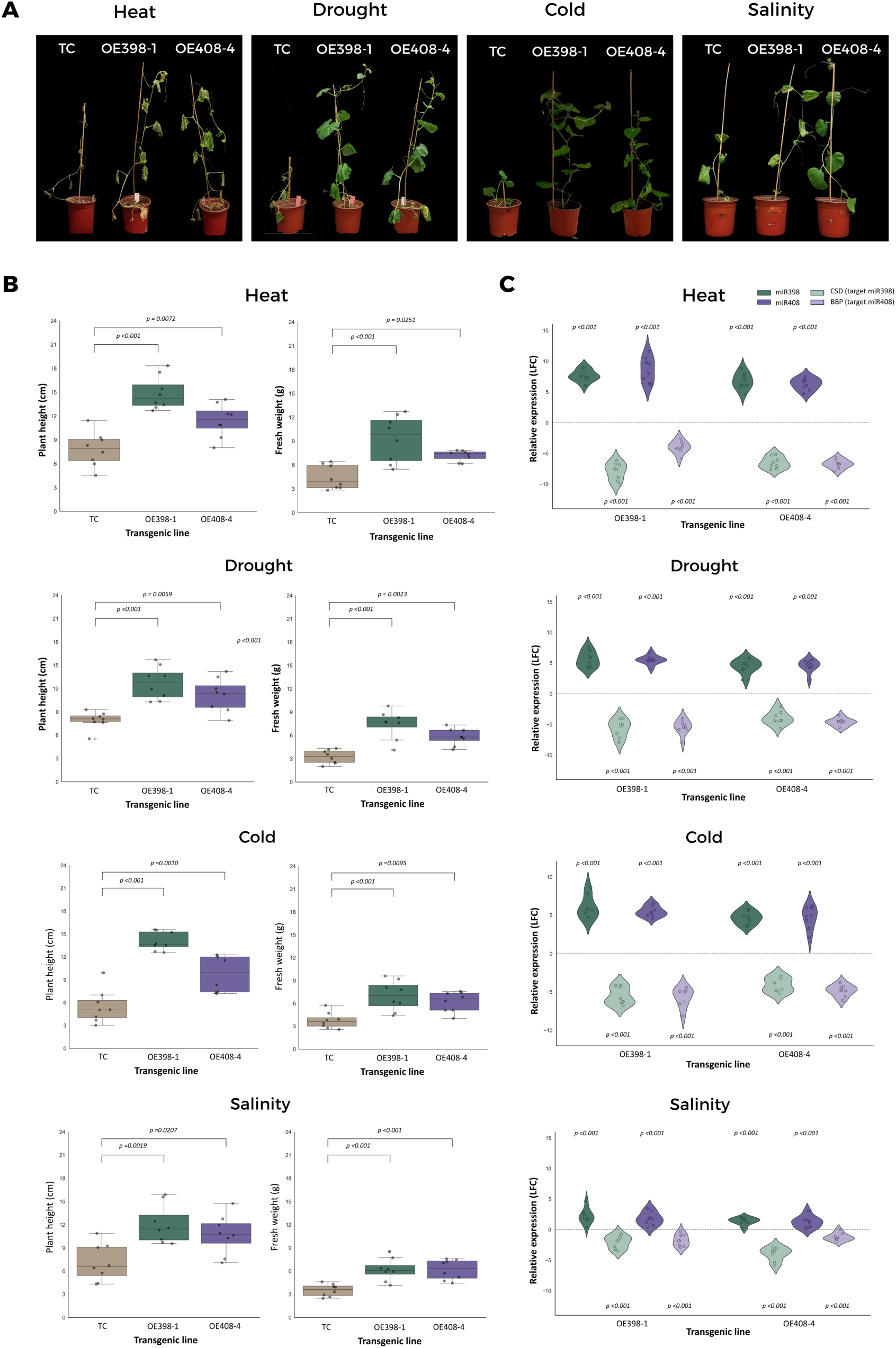
Enhanced tolerance to abiotic stress conditions is associated with the functional activity of miR398a and miR408. A) Representative images of TC, OE398-1, and OE408-4 plants exposed to heat, drought, cold, and salinity for 8 days. B) Box plots quantifying the increased tolerance of OE398-1 and OE408-4 plants under stress conditions, estimated by plant height (left) and fresh weight (right). Differential parameters are represented by the median (internal box line) of the measured values. C) Violin plots showing accumulation levels (estimated by RT-qPCR) of miR398a and miR408 and their respective targets in transgenic plants exposed to the four stress conditions. The miR398a–CSD module is shown in green, and the miR408–BBP module in magenta.

This general observation was reinforced when plant height and biomass production (estimated by fresh weight) were analyzed in details at 8 days post stress treatment (Fig. 6B). The values of both stress-tolerance indicators (plant height and biomass) were significantly higher in the transgenic lines, showing a 1.5- to 2-fold increase in median values compared to TC plants under the four stress conditions. The most pronounced differences were observed under cold-induced stress, with median plant heights ranging from 11.5 to 13 cm in OE398-1 and OE408-4 lines, compared to 4.5 cm in TC plants. Under heat-induced stress, fresh weight values ranged from 7.5 to 10 g in OE398-1 and OE408-4 lines, while TC plants remained below 4 g. In general, OE398-1 plants consistently exhibited higher values than OE408-4 across analyzed parameters (Fig. 6B).

To determine whether the observed stress tolerance was associated with functional activity of the overexpressed miR398a and miR408, we quantified the accumulation levels of both regulatory miRNAs and their target transcripts. As shown in Figure 6C, at 8 days post-treatment, OE398-1 and OE408-4 plants exposed to drought, heat, cold, and salinity exhibited significantly increased miRNA accumulation, accompanied by reduced levels of target transcripts. These differences in stress-tolerance indicators (estimated by plant height) and miRNA/target expression between OE398-1/OE408-4 and TC lines remained relatively consistent throughout the analyzed period (Supplementary Fig. 6).

Considering that drought was the stress condition associated with the highest survival rate, we also analyzed the tolerance of OE398-1, OE408-4, and TC lines to prolonged water deprivation (20 and 35 days), as well as their recovery capacity after irrigation. After extended drought exposure, OE398-1 and OE408-4 plants maintained a more vigorous architecture, characterized by higher leaf density and reduced wilting, whereas TC plants exhibited clear stress symptoms, including chlorosis, loss of turgor, and pronounced size reduction (Fig. 7A and 7B). Upon irrigation after 35 days of water deprivation, OE398-1 and OE408-4 plants showed significantly improved recovery rates (89.7% and 80.15%, respectively) restoring leaf structure and resuming growth, with median biomass values (estimated by fresh weight) of 47 g and 43 g. In contrast, TC plants demonstrated a limited recovery rate (12%) and significantly lower biomass production, with median values of 10 g (Fig. 7C and 7D).

**Figure 7.**
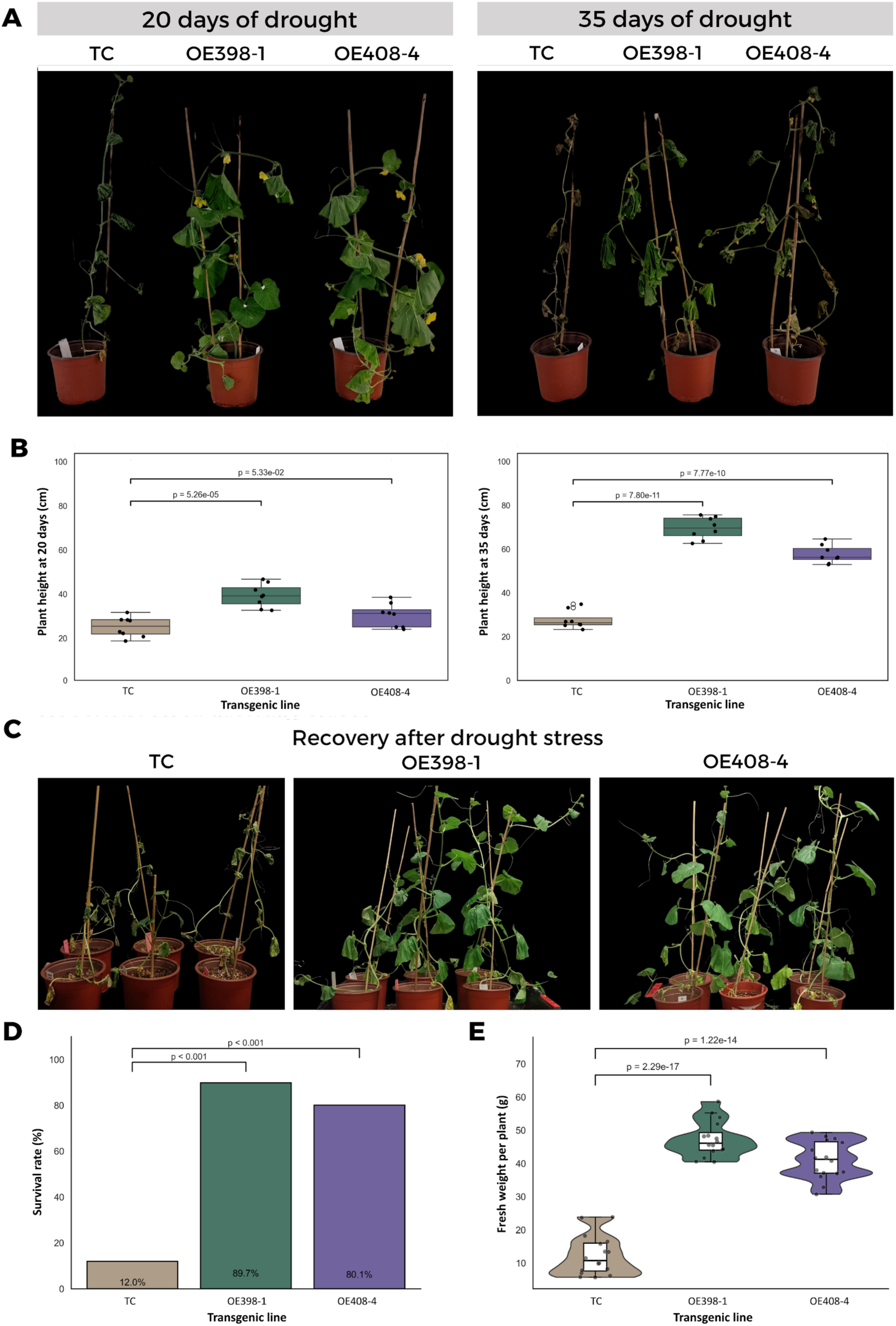
OE398-1 and OE408-4 plants exhibit enhanced recovery after prolonged drought-induced stress. A) Representative images of OE398-1, OE408-4, and TC plants exposed to 20 and 35 days of water deprivation. B) Box plots representing the enhanced tolerance of OE398-1 and OE408-4 plants to prolonged drought conditions, estimated by plant height. C) Representative images of OE398-1, OE408-4, and TC plants at 8 days after irrigation. Graphic representation of the recovery rates of OE398-1 and OE408-4 plants, estimated by survival rate (D) and fresh weight (E).

## DISCUSSION

Climate change poses increasing challenges to agricultural productivity and global food security, primarily due to the rising frequency and intensity of environmental stressors. Without the implementation of innovative biotechnological approaches aimed at enhancing crop tolerance to adverse conditions, significant reductions in crop yields are projected in the near future as a consequence of the climate change scenario (Arif et al., 2025).

Increasing evidence supports the key role of miRNAs in regulating plant responses to stress. Consequently, the fine-tuning of MIR genes or miRNA levels represents a promising biotechnological strategy to enhance stress tolerance in crops (Basso et al., 2019; Raza et al., 2023). Besides more specific functions demonstrated for diverse miRNAs in the regulation of specific response stress, in the last years has been suggested that miR398 and miR408 can act as master regulators of a highly-conserved common response to multiple stress conditions (Sanz-Carbonell et al., 2019, 2020; Villalba-Bermell et al., 2021). Members of the miR408 and miR398 family constitute an ancient and conserved group of miRNAs that predates the divergence of gymnosperms and angiosperms 305 million years ago (Chavez Montes et al., 2014; Li et al., 2022; Gao et al., 2022).

In this study, we demonstrated that overexpression of miR398a and miR408 precursors in melon leads to a significant increase in the accumulation of canonically processed mature miR398a and miR408 forms. Transgenic plants overexpressing miR398a and miR408 exhibited vigorous growth phenotypes, with notable improvements in both vegetative and reproductive traits, including enhanced stem elongation, internode development, more complex root architecture, increased biomass yield, and higher flower production. Upon analyzing their response to environmental changes, we observed that OE398-1 and OE408-4 plants showed improved tolerance to drought, cold, salinity, and heat, reinforcing the proposed key role of miR398 and miR408 in regulating stress responses across diverse plant species (Hang et al., 2021; Hong et al., 2024; Li et al., 2020; Lu et al., 2022a).

Small RNAs sequencing data evidenced that the elevated levels of miR398a in OE398-1 plants were closely associated with a comparable increase in miR408 levels. Similarly, a significant accumulation of miR398a was also detected in the OE408-4 plants, supporting the idea that overexpression of one of these miRNAs exerts a direct positive effect on the accumulation of the other. In parallel, miR398a and miR408 overexpression was also associated with a comparable increasing in miR397 accumulation in transgenic plants, suggesting the existence of a coordinated and interdependent strategy for the expression of these miRNAs in melon. It is important to note that miR398a, miR408 and miR397 form part of a group of riboregulators know as copper miRNAs (Cu-miRNAs), that are predominantly involved in the regulation of copper protein expression in plants (Burkhead et al., 2009; Pilon, 2017). Although previous reports have shown that the transcription factor SQUAMOSA Promoter binding protein-Like7 (SPL7) acts as a common positive regulator of Cu-miRNAs in plants (Yamasaki et al., 2009; Li et al., 2022; Yang et al., 2024), the miRNA-dependent co-regulation between Cu-miRNA observed in melon has not been described in other species. However, it is important to note that in several previous studies analyzing the effects of constitutive overexpression of individual Cu-miRNAs, their impact on the overall small RNA (sRNA) population in transgenic plants has not been evaluated (Hong et al., 2024; Yang et al., 2024; Song et al., 2018; Lu et al., 2022a). Further studies are needed to elucidate the molecular basis and broader relevance of this potential co-regulatory phenomenon.

As expected, coordinated overexpression of Cu-miRNAs was functionally associated with the significant down regulation of their well-established targets in melon, CSD (target of miR398a), BBP (target of miR408) and a Laccase-related gene (targets of miR397). Consequently, is pertinent to assume that, in coincidence with the observed in arabidopsis (Song et al., 2018), poplar (Guo et al., 2023), tobacco and rice (Pan et al., 2018), miR408 and/or miR397 over expression may be involved in the vigorous phenotype observed in OE398-1 and OE408-4 plants. In these species, has been demonstrated that miR397-mediated laccase down regulation reduces excessive lignification, resulting in more flexible tissues, enhanced stem elongation and improved root development (Hong et al., 2024). In parallel, miR408 regulate negatively copper-binding proteins (BBP, in melon) enhancing photosynthetic efficiency contributing to the plant biomass increasing (Yang et al., 2024).

A broader analysis of the molecular functions in the shared DEG in both transgenic lines revealed a predominant decreasing in Cu⁺-binding activity, directly associated to the overaccumulation of the cu-miRNAs (miR408, miR398a and miR397). As mentioned above, these miRNAs were broadly associated to the regulation of the stress response in diverse plant species (Lyu, 2024). In a recent study, demonstrating that the simultaneous overexpression of three Cu-miRNAs (miR397, miR408 and miR528) induce enhanced tolerance against cold and drought in rice and maize, was suggested that stress-resistance is associated to silencing of laccase (LAC) genes that diverts metabolism from lignin synthesis to flavonoid accumulation (Hong et al., 2024). In contrast, in arabidopsis plants in that miR408 expression was suppressed, was proposed that the enhanced drought-tolerance observed in these plants was associated to increased accumulation of the miR408-target blue copper protein PLANTACYANIN (PCY), promoting ROS accumulation and stomatal closure (Yang et al., 2024). Regarding the highly conserved miR398-CSD module, an inconsistent response has been described alternating increased/decreased accumulation according to plant specie or stress type analyzed (Li et al., 2022; Zhu et al., 2011). Thus evidencing that although conserved cu-miRNAs compromises stress tolerance in multiple plants, the cellular mechanism underlying its function remains largely unclear and may go beyond the regulation of secondary metabolism (Akhtar et al., 2025).

Having analyzed as the overexpression of miR398a and miR408 alter the general miRNA and transcriptional landscape of OE398-1 and OE408-4 plants we were able to identify the coordinated and interdependent expression of Cu-miRNAs, evidencing that specific miRNA-target modules analyzed in previous similar works can represent only a part of the a more complex regulatory phenomenon linking Cu-miRNAs and stress response in plants.

It is important to consider that the predominant role of highly conserved cu-miRNAs is regulate the cu-homeostasis in the plant trough the fine modulation of the Cu availability (Akhtar et al., 2025; Pilon, 2017). However, Cu deficiency is a rare event in most natural soils (Burkhead et al., 2009), arguing against the idea that these miRNAs may have evolved exclusively as an adaptation to Cu-limited environments (Pilon, 2017).

It has been proposed that cu-miRNAs may act as mobile signals able to modulate at local and systemic level the distribution of Cu in the cell allowing plants to coordinate development and response to stress (Pilon, 2017). The highly co-dependent expression observed for miR398a, miR408 and miR397 and their copper-related targets plus the significant increased-expression of melon genes with molecular functions related to transmembrane transporter activity observed in OE398-1 and OE408-4 plants, reinforce this possibility.

Overall, our study offers a comprehensive analysis of the effects of the overexpression of miR398 and miR408 (two members of the Cu-miRNA family) and the resultant increased tolerance to diverse abiotic stress conditions such as drought, cold, salinity and heat in melon plants. Considering that Cu-miRNAs constitute are evolutionary conserved in plants, this bioengineering approach could constitute a valuable and broadly applicable strategy for developing plants with enhanced adaptability to diverse adverse environmental conditions.

## METHODS

### Plasmid construction

Pre-miR398a (100 bp) and pre-miR408 (134 bp) precursors were amplified using oligonucleotide primers (MIR398a-F and MIR398a-R or MIR408-F and MIR408-R) containing recognition sites for BsaI restriction endonucleases. The PCR products were BsaI-digested and ligated into the binary vector pMCD32, that contain a ccdB gene and a chloramphenicol resistance (CmR) cassette flanked by BsaI sites to allow…. All constructs included the phosphinothricin acetyltransferase (bar) gene, which confers resistance to the herbicide glufosinate, an inhibitor of (RS)-2-amino-4-(hydroxy(methyl)phosphonyl) butanoyl) amino acid synthesis (Becker et al., 1992; Thompson et al., 1987), under the control of the NOS promoter and its enhancer. The binary vectors pMCD32-miR398a and pMCD32-miR408, containing the miR398a and miR408 precursor gene cassette, respectively, were transferred into *Agrobacterium tumefaciens* strain AGL0 by electroporation [15]. An empty vector was used to generate transgenic control plants.

### Melon transformation

For melon (*Cucumis melo* cv. Vedrantais) transformation, the technique of cotyledon transformation with *Agrobacterium tumefasciens* strain AGL0 was used, following the previously described method (García-Almodóvar et al., 2017).. Briefly, half of the proximal portions of cotyledons were cut from 1-day-old seeds, which were then co-cultured with transformed *Agrobacterium* for 20 min in the presence of 200 µM acetosyringone. Inoculated explants were co-cultured at 28°C for three days on regeneration medium containing 0.5 mg/l 6-benzylaminopurine (BA), 0.1 mg/l indole-3-acetic acid (IAA) and 200 µM acetosyringone. Every 2 weeks, the green cluster-shaped shoots were cut, and the explants were transferred to fresh selection medium containing L-Phosphinothricin (PPT). Once the regenerated shoots reached a considerable size, they were cut, separated from the explants, and placed individually on rooting media inside large test tubes. When the rooted seedlings reached a suitable size, a leaf was cut to identify the transformed plants.

Genomic DNA was extracted from young leaves of melon plants using an improved cetyltrimethylammonium bromide (CTAB) method. For genotyping of the candidate plants, a region (221 bp) of the bar gene was amplified to confirm the stable transformation. Primers used in this assay are listed in Supplementary Table 1. The ploidy of the transformed plants was determined by flow cytometry (Central Support Service for Experimental Research -SCSIE- Universitat de València). Ten diploid lines, five transformed with the miR398a-vector (OE398-1 to -5), four with the miR408-vector (OE408-1 to -4) and one with the control empty vector (TC), were recovered and selected for generation of stable homozygous plants.

Transformed plants were transferred to the greenhouse facilities of the Polytechnic University of Valencia (Valencia, Spain) and the Center for Research in Agricultural Genomics (CRAG) (Barcelona, Spain), where they were maintained under controlled conditions. During flowering, manual pollination was carried out until fruits were obtained. Plants were purified by two consecutive generations of backcrossing to ensure that each line was homozygous.

### Phenotypic characterization

Seeds of the ten homozygous lines were germinated in Petri dishes at 37°C for 48 h in darkness, followed by 24 h at 25°C with a 16 h light and 8 h dark cycle. Melon seedlings (16 for each line) were planted in pots, maintained under controlled conditions (28°C with 16 h of light and 20°C with 8 h of darkness) and monitored during 80 days.

Plant height, number and length of internodes, total length of the main root, number of lateral roots, length of the taproot, number of leaves and number of flowers per plant were analyzed. Plant height was considered as the distance from the substrate surface to the apex of the main stem. The number of internodes was determined by counting the segments from the cotyledon to the end of the main stem, considering as the first internode the one between the cotyledons and the first true leaf. The length of each internode was measured individually. Root length was determined from the base of the stem to the tip of the main root. In addition, the total number of fully developed leaves and the number of visible flowers per plant at the time of sampling were quantified.

### Stress treatments

Seeds of two representative lines (OE408-4 and OE398-1) selected according to the highest expression levels of their corresponding miRNA-precursors and the transformed control were germinates as described above. Only plants were selected for stress treatments.

At 11 days after emergence, plants with homogeneous development were subjected to the following treatments: 1) Salinity (irrigation with 50 mL of 200 mM LiCl); 2) Cold (irrigation with 50 mL of Hoagland solution and maintained at 20°C/16 h light and 14°C/8 h dark; 3) Drought, (initial irrigation with 50 mL of Hoagland solution and deprivation of irrigation until end of the assay; and 4) Heat (initial irrigation with 50 mL of Hoagland solution and maintained at 42°C/16 h light and 32°C/8 h dark. Control transformed plants were irrigated with 50 mL of Hoagland’s solution. Three replicates per treatment were performed, and each biological replicate analyzed corresponded to a pool of eight plants. Except for the drought treatment, plants were flood irrigated by alternating water and Hoagland solution (1500 mL every 48 h); except for the cold treatment, plants were maintained at 28°C/16 h light and 20°C/8 h dark. At 11 days post-treatment, when all plants had six true leaves, the first leaf below the apex of each plant was collected, frozen in liquid nitrogen and stored at -80°C until analysis. This plant material was used as a source for all analyses performed in the present study.

### RNA analysis

Total RNA was extracted from melon plants using TRIzol extraction protocol (Thermo Scientific ™). RNA quality was assessed by electrophoresis using 1% (w/v) agarose gel, followed by spectrophotometric quantification. cDNA was synthesized from 1 ug of total RNA using the RevertAid cDNA Synthesis Kit with dsDNAse (Thermo Scientific ™). For real time qPCR, 30 ng of cDNA were analyzed using real-time PCR QuantStudio Real-Time PCR instrument (Thermo Scientific™). PyroTaq EvaGreen mix Plus (ROX; CulteK Molecular Bioline, Madrid, Spain) was used to monitor dsDNA synthesis in accordance with the manufacturer’s instructions. Expression of ADP-ribosylation factor (MELO3C018705) and Profilin (NM_001297545.1) were used as the housekeeping gene for internal normalization.

Small RNAs were isolated by using REALTOTAL microRNA Kit RBMER14 (Durviz, Paterna, Valencia, Spain) according to the manufacturer’s instructions, and quantified using the stem-loop RT-PCR technique (Kramer, 2011). cDNA was synthesized using RevertAid First Strand cDNA Synthesis Kit (Thermo Scientific™). qPCR amplification was performed on 50 ng of cDNA, as described above. Normalization was performed by using U6 snRNA (MELO3C005454) as internal control. Table S10 gives a list of the primers used for qPCR analysis.

### Small RNA Sequencing and Data Analysis

A quality control of the small RNA (sRNA) reads was performed using FastQC (v0.12.1) (Andrews, 2019) and the raw sequencing data were adapter- and quality-trimmed using fastp (v0.24.0) (Chen, 2023), retaining only reads between 20 and 25 nucleotides in length. All retained reads were aligned with Bowtie (v1.3.1) (Langmead et al., 2009), with --best -v0 -k1 parameters, to the precursor regions corresponding to the mature miRNA of MIR398a and MIR408 genes. The mean of reads per million (RPMs) was calculated for each unique read within each sample group (OE398-1, OE408-4 and control).

The accumulation profile of the different sRNA size was analized among group using a Bray–Curtis distance matrix. Differences in overall composition were assessed with PERMANOVA using 9,999 unrestricted permutations. Then, the reads were mapped to ribosomal RNA (rRNA) sequences from the Viridiplantae clade in RNAcentral (The RNAcentral Consortium, 2019) using Bowtie with the same parameters. This step aimed to remove rRNA from the libraries.

An absolute count matrix was generated from the remaining reads. Low-abundance sequences, defined as those with fewer than five counts in at least three samples, were filtered. The result matrix was normalized by size factor and transformed with the vst function, with the “blind” option enabled, from DESeq2 (v1.38.3) (Love et al., 2014) in R (v4.2.0). Principal component analysis (PCA) was then performed by selecting the 2000 features with the highest variance using the prcomp function of the stats R package and it was visualized by ggplot2 (v3.5.1). Sample coordinates on the first three principal component (PC1-PC3) were used to compute pairwaise Euclidean distances. A one-side Mann-Whitney-Wilcoxon test was applied to assess wheter between-group distances were significantly (p-value > 0.05) greater than within-group distances.

For differential expression analysis of small RNAs (sRNAs), DESeq2 R package was used. Normalization of raw counts was performed using the DESeq2 median of ratios method. Hypothesis testing was performed by a Wald test, and p-values were adjusted for multiple testing using the Benjamini–Hochberg (BH) procedure. The lfcShrink() function in DESeq2 was applied to shrink fold changes. Sequences were considered differentially expressed at an adjusted p-value threshold of <0.05.

The annotation of miRNAs was performed by mapping with Bowtie2 (v2.4.4) (Langmead and Salzberg, 2012) (parameters: --local -N0 -L20 –score-min C,20,0), the mature miRNAs of *Cucumis melo* in miRBase (https://www.mirbase.org). Identified miRNAs were analyzed at the family level, disregarding subfamily classifications. For miRNA families represented by more than one sequence, the median LFC of the individual sequences was used as the representative value for that family. Analysis data are presented in Table S11.

The remaining sRNAs were annotated by aligning them with Bowtie (parameters --best - v2 -k1) against sequences classified as sncRNAs in RNAcentral and not aligned sRNAs were aligned to the *Cucumis melo* (DHL92 v4) genome, obtained from the Cucurbit Genomics Database v2.0 (http://cucurbitgenomics.org/v2/) (Yu et al., 2023). The genome alignment was performed by Bowtie with the same parameters as RNAcentral. When a given sRNA sequence matched multiple genomic features, annotations were assigned following the priority order: five_prime_UTR > three_prime_UTR > CDS > exon > gene (Intronic region) > miRNA precursor. Finally, sequences that did not align or that did not correspond to any annotated genomic region were classified as other sRNAs.

### Transcriptome Sequencing and Data Analysis

A quality control step was performed for the RNA-Seq (mRNA) reads using FastQC. The raw sequencing data were adapter- and quality-trimmed using fastp. Transcript quantification was performed using the pseudo-alignment tool Salmon (v1.6.0) (Patro et al., 2017). As a reference, *Cucumis melo* (DHL92 v4) genome.

Salmon results were imported using the tximport R package (v1.26.1) (Soneson et al., 2016) with a tx2gene file to aggregate the counts to the gene level. Principal component analysis (PCA) and differential expression analysis were performed following the same procedures as sRNA analysis. The only difference in the pipeline was the low-abundance filtering step, where genes with fewer than five counts in at least two samples were removed, and differential expression was determined with a threshold of adjusted p-value <0.05 and a LFC > |0.580|.

The relationship between miRNAs and their targets was inferred from a file containing validated miRNA–target gene modules in *Arabidopsis thaliana*, obtained from TarDB (http://www.biosequencing.cn/TarDB/). BioMart of Ensembl plants (https://plants.ensembl.org) was then used to identify the *Cucumis melo* orthologues of the target genes. Analysis data are presented in Table S12.

### Gene Ontology Enrichment

Gene Ontology (GO) enrichment analysis was performed using the clusterProfiler package (v4.6.2) (https://doi.org/10.1016/j.xinn.2021.100141), with the org.At.tair.db annotation database (v3.16.0) as the reference. Since a GO annotation database is not currently available for C. melo, all differentially expressed genes (DEGs) identified in melon were first mapped to their orthologs in A. thaliana using an orthology table downloaded from BioMart (Ensembl Plants). The entire set of *A. thaliana* genes with an identified ortholog in *C. melo* was used as the background. The analysis was conducted within the Biological Process and Molecular Function categories, including terms associated with gene sets ranging from a minimum of 10 genes to a maximum of 500 genes. A multiple testing correction was applied using the BH method, with a significance threshold set at an adjusted p-value < 0.05. To reduce redundancy, the Wang method was applied with a redundancy threshold of 0.7. This analysis was performed for commonly upregulated and downregulated DEGs between OE398-1_vs_control and OE408-4_vs_control contrasts.

