## Supplementary material for "Coordinated overexpression of miR398 and miR408 confers broad-spectrum abiotic stress tolerance in melon": Figure Sup

**Figures Supplementary:**

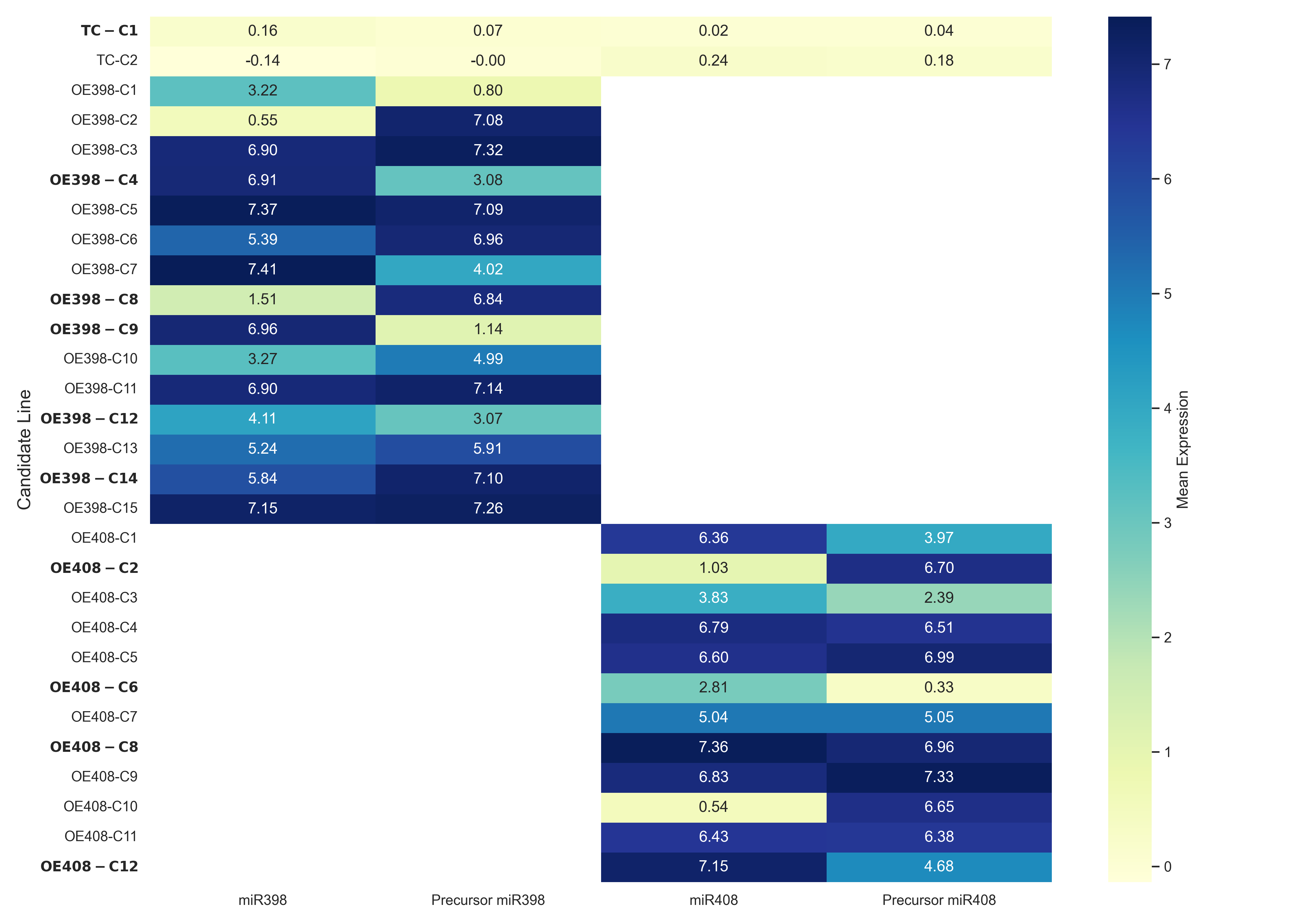

Figure S1. Detailed description of the transformed plants selected for ploidy analysis. Accumulation values (estimated by stem-loop RT-qPCR and conventional RT-qPCR) of miR398a and its respective precursors (left) and miR408 and its respective precursors (right) are shown. Diploid candidate plants marked in bold were selected based on high miRNA/precursor accumulation and renamed as follows for subsequent studies: TC-C1 (TC), OE398-C4 (OE398-1), OE398-C8 (OE398-2), OE398-C9 (OE398-3), OE398-C12 (OE398-4), OE398-C14 (OE398-5), OE408-C2 (OE408-1), OE408-C6 (OE408-6), OE408-C8 (OE408-3), and OE408-C12 (OE408-4).

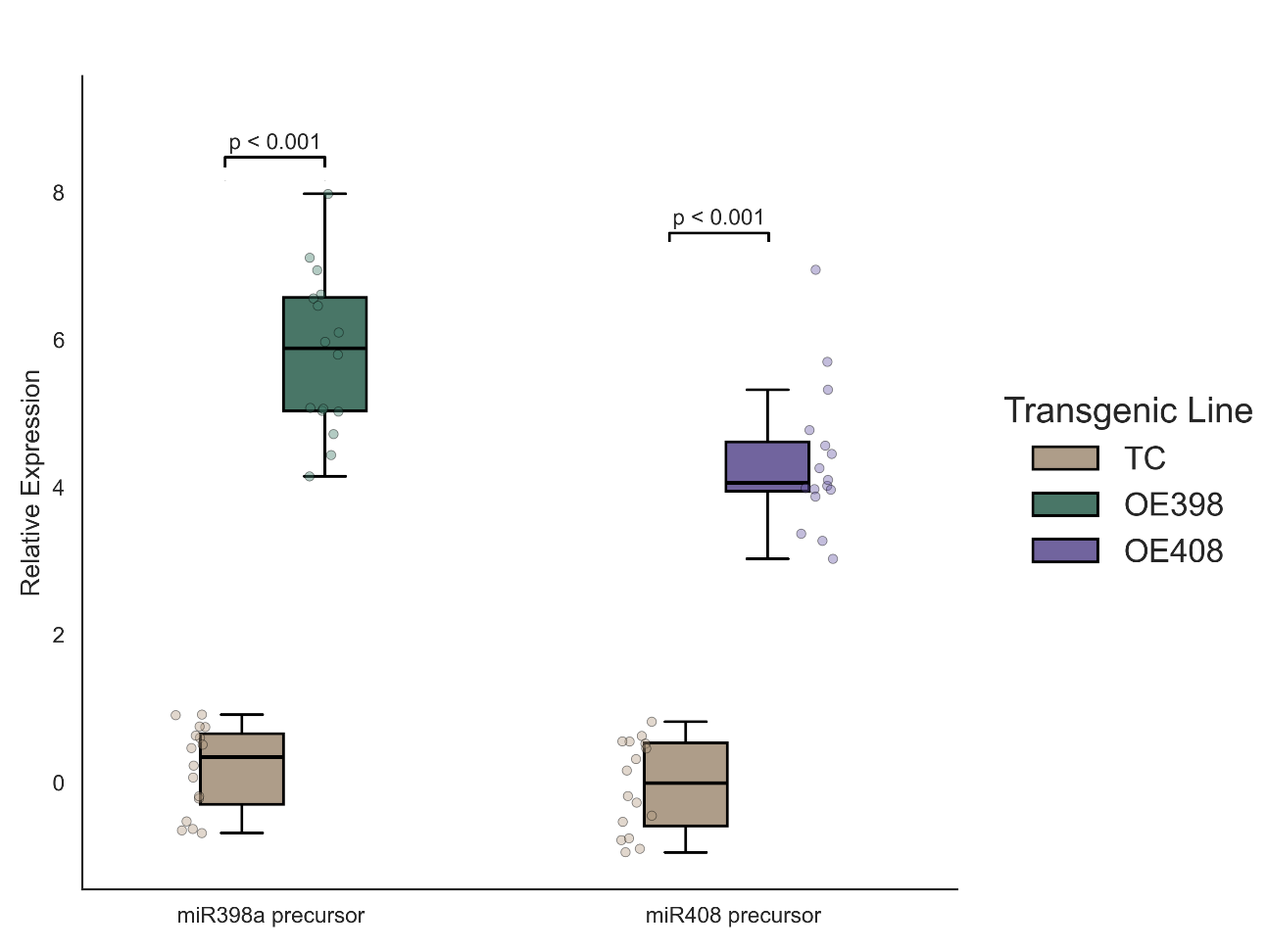

Figure S2. Plants of OE398-1 and OE408-4 lines exhibit increased accumulation of miRNA precursors. Box plots showing the significant increase in the accumulation of the miR398a precursor and the miR408 precursor in OE398-1 and OE408-4 plants, respectively. Dots (green for miR398 precursors and magenta for miR408 precursors) represent values obtained from 15 individual plants analyzed for each transgenic line.

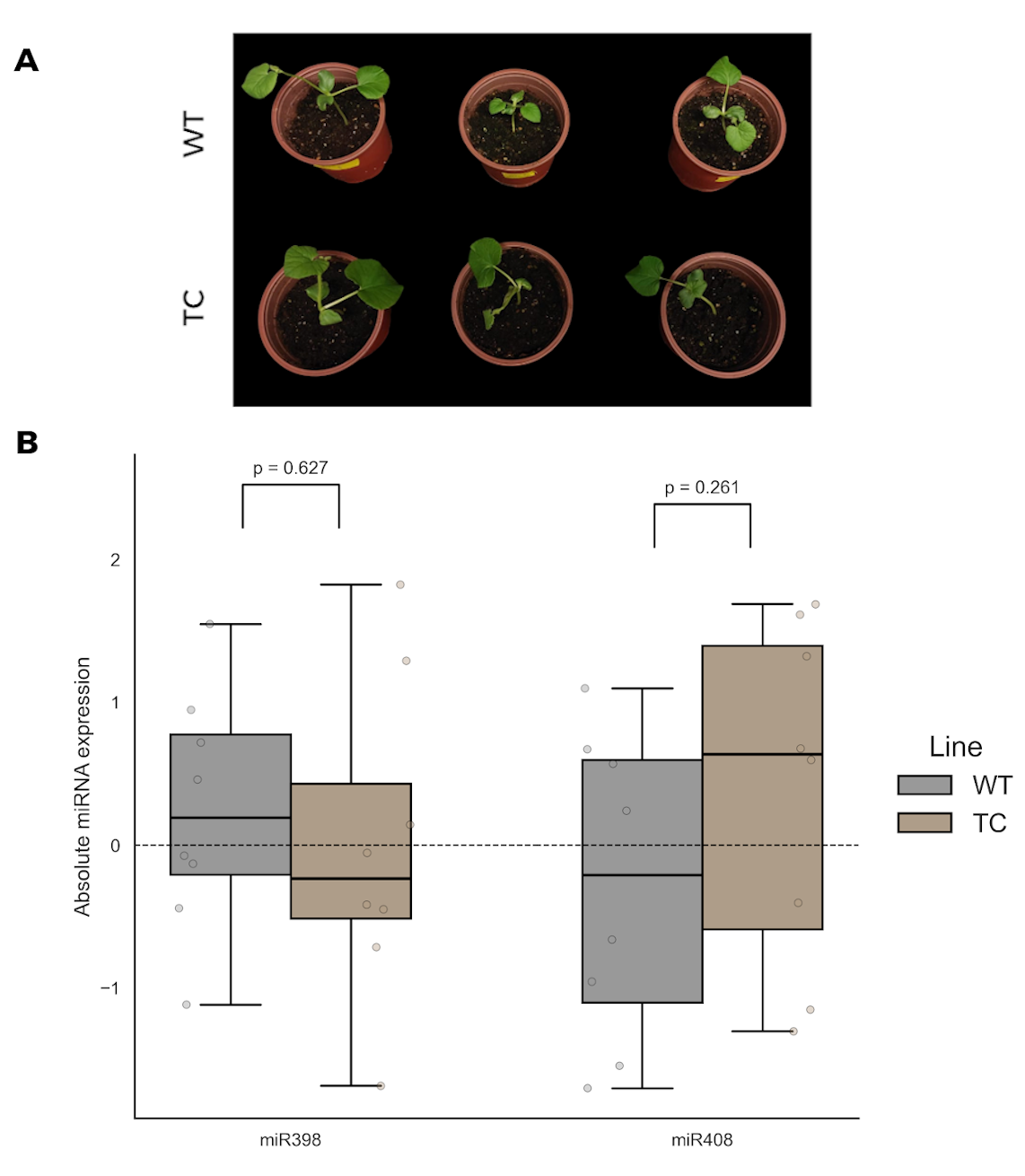

Figure S3. Untransformed (WT) and transformed control (TC) plants exhibit comparable characteristics. A) Image of three representative WT and TC plants at 4 days post emergence. B) Box-plot showing the comparable accumulation of the miR398a and miR408 precursors in WT and TC plants. Dots (gray for WT and brown for TC) represent values obtained from 8 individual plants analyzed for each group.

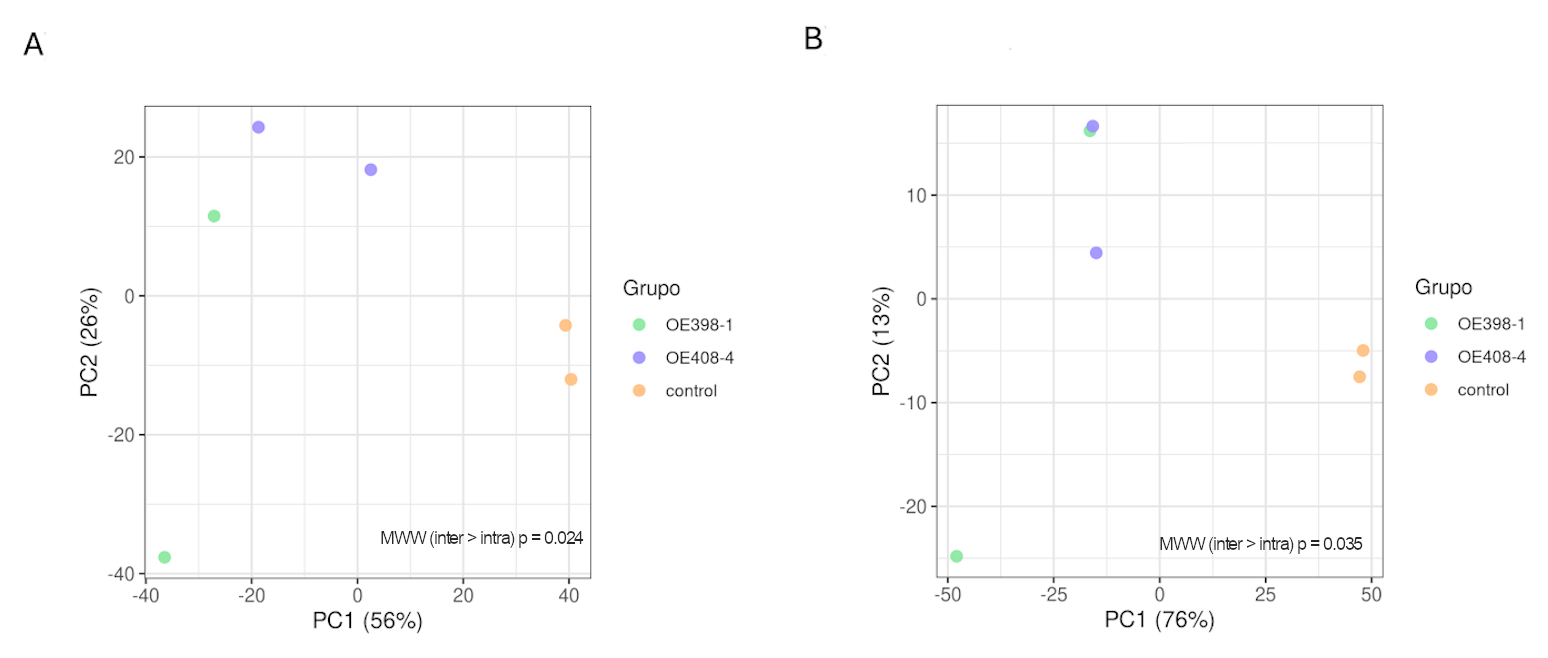

Figure S4. Principal component analysis of sRNA sequences (A) and transcripts (B) recovered from OE 398-1, OE408-4 and TC plants. Values of proportion of variance for PC1 and PC2 are showed in the X- and Y-axis. The statistical significance was estimated by Mann-Whitney-Wilcoxon test, considering the inter- and intra-group distances (*P*-values are showed in the graphic).

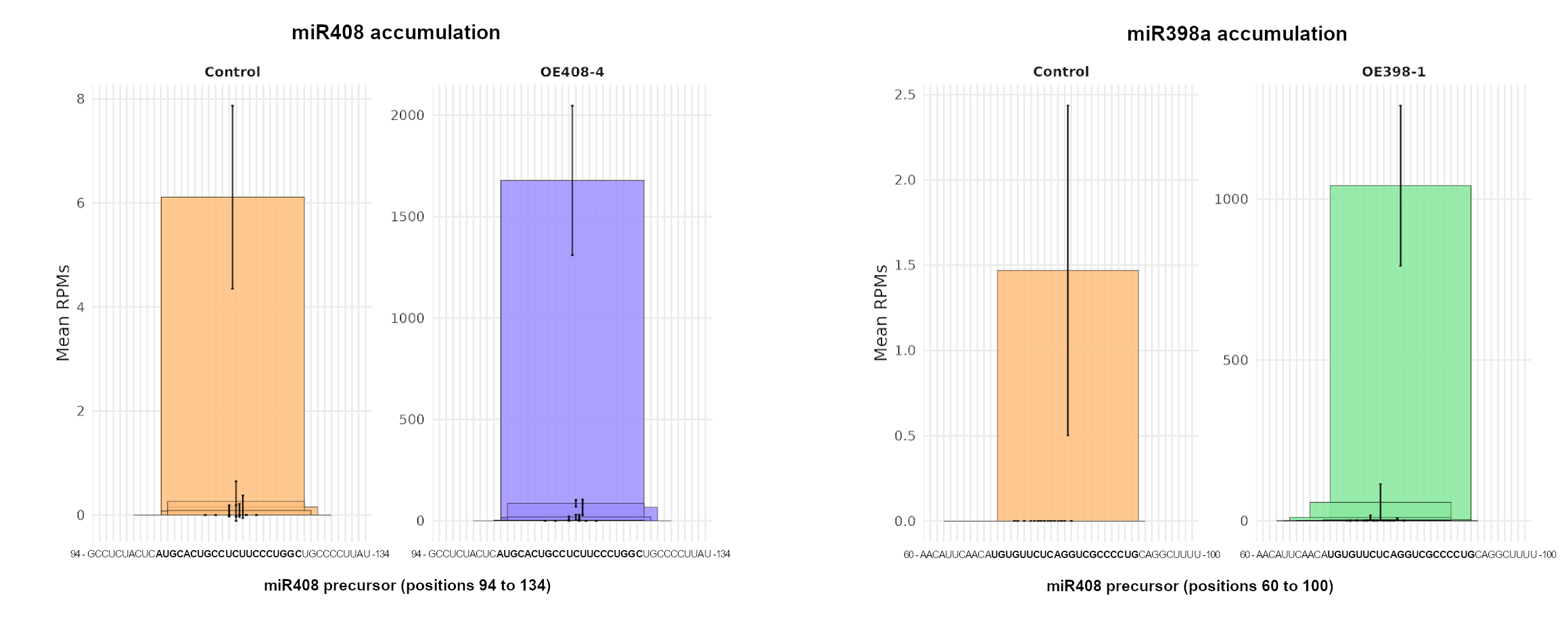

Figure S5. The miR398a and miR408 overexpressed precursors are accurately processed in transgenic plants. Graphic representation of miR408 (left) and miR398a (right) sequences recovered from the sRNA dataset. The predominant overaccumulated sequences correspond to the canonical mature forms of miR398a (green) and miR408 (magenta), highlighted in bold within their respective precursor sequences. The corresponding miRNA sequences recovered from TC plants are shown in orange. Values represent the mean RPMs of sequences per library.

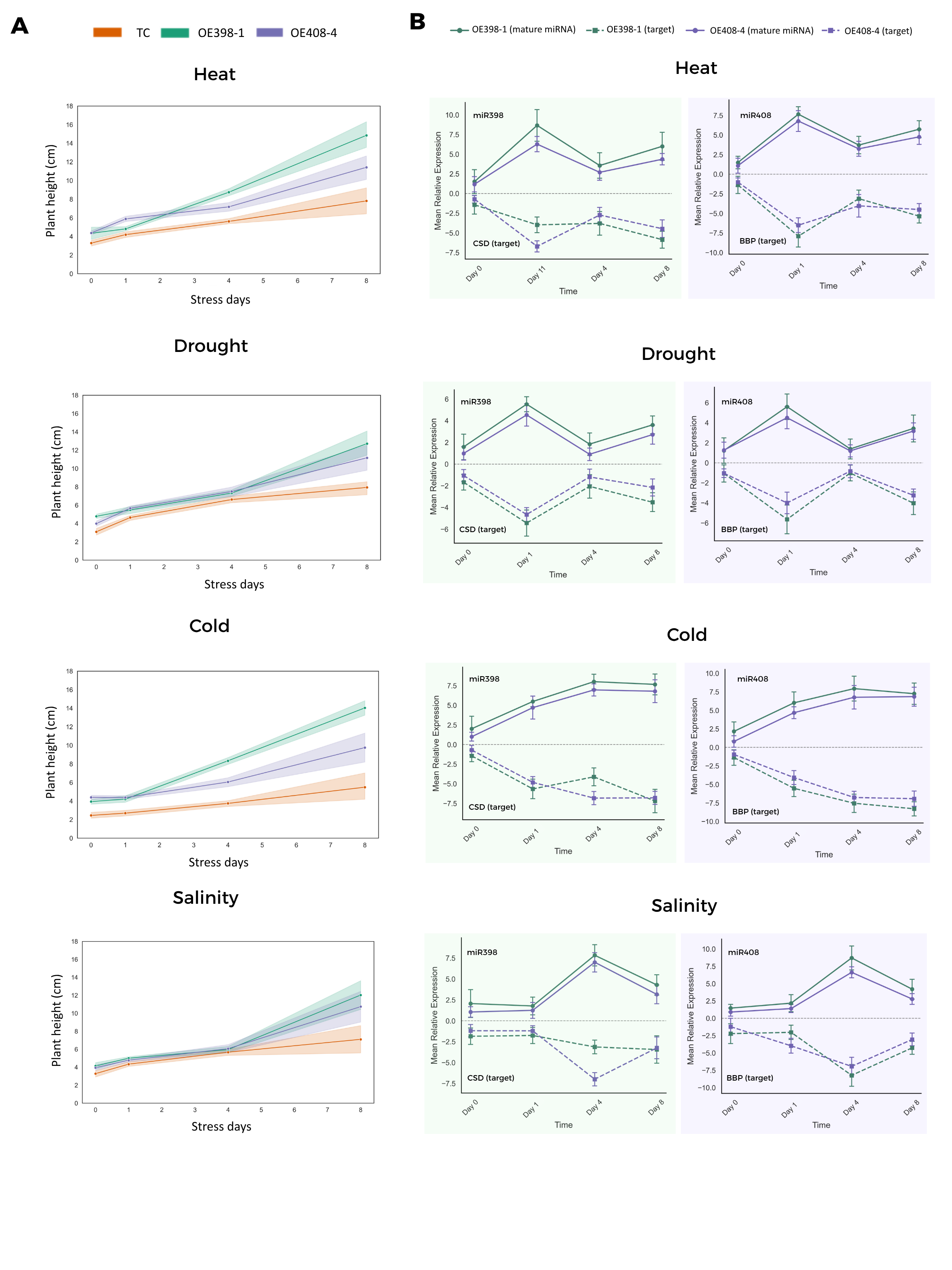

Figure S6. Stress tolerance observed in OE398-1 (green lines) and OE408-4 (magenta lines) plants (left panels) is associated with the functional activity of the regulatory modules miR398a-CSD (green panels) and miR408-BBP (magenta panels). Stress tolerance was estimated by plant height. Accumulation of miRNAs and their target transcripts was assessed using stem-loop RT-qPCR and conventional RT-qPCR, respectively.
